# abCAN: a Practical and Novel Attention Network for Predicting Mutant Antibody Affinity

**DOI:** 10.1101/2024.12.02.625958

**Authors:** Chen Gong, Yunyao Shen, Hongde Liu, Wenlong Ming

## Abstract

Accurate prediction of mutation effects on antibody-antigen interactions is critical for antibody engineering and drug design. In this study, we present abCAN, a practical and novel attention network designed to predict changes in binding affinity caused by mutations. abCAN requires only the pre-mutant antibody-antigen complex structure and mutation information to perform its predictions. abCAN introduces an innovative approach, Progressive Encoding, which progressively integrates structural, residue-level, and sequential information to construct the complex representation in a systematic manner, effectively capturing both the topological features of the structure and contextual features of the sequence. During which, extra weight to interface residues would also be applied through attention mechanisms. These learned representations are then transferred to a predictor that estimates changes in antibody-antigen binding affinity induced by mutations. On the benchmark dataset, abCAN achieved a root-mean-square error (RMSE) of 1.195 (kcal/mol^-1^) and a Pearson correlation coefficient (PCC) of 0.841, setting a new state-of-the-art (SOTA) benchmark for prediction accuracy in the field of antibody affinity prediction.

## 1 Introduction

Antibodies, which are multimeric proteins with molecular-specific recognition capabilities to elicit immune responses [1], play an essential role in the human immune system. Antibody-antigen interaction occurs between the complementary determining regions (CDRs) of the antibody and a specific epitope on the antigen, making it specific and selective [2]. Antibodies have demonstrated superior performance over traditional treatments for certain cancers, with fewer side effects and lower resistance [3]. Antibody drugs have consistently topped the global bestseller lists for years and represent the fastest-growing segment in the pharmaceutical industry [4].

Antibody optimization is the core step of developing novel antibody drugs, which focuses on identifying antibodies with higher binding affinity while exhibiting drug-like properties through mutation or engineering [5]. Binding free energy (Δ*G*) is commonly used to represent binding affinity, as it reflects the thermodynamics of antibody-antigen interactions [6]. Currently, developing antibody drugs still relies on labor-intensive in-vitro techniques to screen for mutant antibodies with higher binding affinity [7], which are time-consuming and labor-intensive. Consequently, in-silico tools capable of evaluating affinity changes upon mutations (i.e.,ΔΔ*G*) are in urgent need of rapid screen for high-affinity antibodies [8].

In recent years, three types of computational approaches have emerged [9]. The first is molecular modeling approaches, which would coarse-grained amino acid molecules to simulate the dynamics of antibody interactions, offering reliable results but at the cost of extremely high computational demands [10]. The second is energy function-based algorithms. Traditional energy function-based methods utilize proven biological mechanism and statistical probability [11], scoring the binding interactions between target antibodies and antigens, with these scores serving as the criteria for affinity determination. For instance, the CUMAb method proposed by Tennenhouse et al. combines structural information and considers both energy and structural integrity to identify optimal antibodies [12]. The major issue with these methods is their lack of accuracy, since the design of energy functions is based on empirical considerations [13]. The interactions of macromolecular proteins at the microscopic level are not fully understood yet, significantly limiting the potential for further improvements in algorithm accuracy.

The last category is machine learning-based prediction models. Machine learning paradigms, particularly deep learning, have gained popularity during the past years [14]. The accumulated experimental data and existing knowledge relating to antibody has provided an unprecedented opportunity for these methods. Machine learning methods can implicitly learn the interaction rules between antibodies and antigens [15], replacing the manual design process of energy functions. Studies have shown that unsupervised learning can capture intrinsic properties of antibodies, such as stability, from extensive and diverse protein sequence databases. Supervised learning, on the other hand, can learn specific exogenous properties, such as antibody affinity, using experimental data on antibody-antigen interactions [16].

The main limitation lies in the scarcity of structural data [17], combined with the effective representation and utilization of limited data [18]. Two approaches currently address these issues:

Firstly, leveraging sequence data that is available in large quantities [19]. Some studies rely solely on antibody sequences to build models, or fine-tune pre-trained protein language models for downstream antibody sequence prediction tasks [20]. For instance, Li et al. demonstrated that models pre-trained with millions of human antibody sequences achieved better performance [21]. However, we cannot exclude the use of structural data, as affinity is closely related to the three-dimensional binding conformation of antibody epitopes and antigenic determinants [22]. Jin et al. developed the antibody design model HSRN, which integrates structural information into training sequences, and its superior performance underscores the biological importance of structural data [23].

Another approach focuses on designing better representations, maximizing the use of limited data and enabling the model to learn essential features from the raw data [24]. Miller et al. found that focusing on simple features of antibody-antigen binding information can achieve the same learning effectiveness as complex features involving energy, statistics, and networks [25]. Therefore, emphasizing contact surface representations rich in biological interaction information could play a crucial role in breaking through model performance barriers [2].

Herein, based on previous solutions, we developed abCAN—Antibody-Antigen Complex Attention Network—a practical and novel architecture for predicting antibody affinity. abCAN has an encoder for constructing antibody-antigen complex representation and a predictor for predicting ΔΔ*G*. The encoder comprises two multi-layer message passing networks (MPN) modules, integrated with bidirectional gated cycle unit (GRU) and attention mechanisms. We innovatively proposed an encoding architecture, Progressive Encoding, which leverages the two MPN modules to separately learn the geometric features of the complex structure and the intrinsic properties of amino acids, while the GRU captures the contextual information of complex sequence and incorporates it into the overall representation through attention mechanisms. abCAN predicts changes in affinity values for mutated antibodies based on known antibody-antigen complexes. We evaluated abCAN’s predictive ability on a widely used benchmark dataset, assessing its accuracy and robustness through statistical methods and cross-validations. abCAN achieves new SOTA performance compared to existing advanced models. This feature integrating approach to data representation effectively addresses the issue of scarce structural data, endowing our model with higher accuracy.

## 2 Materials and methods

### 2.1 Dataset Construction and Training Set Augmentation

We collected data for model training and evaluation from three datasets: AB-BIND, SKEMPI2.0 and SKEMPI [26][27][28]. For duplicate data in these databases, we referred to the screening criteria from mCSM-AB2 [29], retaining data where the experimental method was SPR or ITC and the experimental information was complete.

As subsets of these datasets, S1131 and M1707 are two benchmark datasets widely used to test the predictive performance of protein-protein interaction models [30]. We selected the type of antibody-antigen interaction within them as benchmark dataset of this work and named it M305, which contains a total of 305 data containing both single and multiple mutations. This benchmark dataset is used to compare the performance of abCAN with other SOTA models.

Besides data in the benchmark M305, the remaining data within the three source datasets is then divided in a ratio of 4:1 into training set and validation set. Each data is comprised of the change in affinity (ΔΔ*G*) caused by mutations, the pre-mutant antibody-antigen complex PDB file, and the mutation information (**Figure 1a**). Among the training set, the majority ΔΔ*G* values are negative [8], displaying a skewed distribution. To balance it, we introduced hypothetical reverse mutations (**Figure 1b**) [31]. For training data with ΔΔ*G* values within the range of [-2, 2] [32], we added a reverse version of it to the training set, with inverted wild-type and mutant, as well as a negated ΔΔ*G* values. This approach also serves as a method of data augmentation [33]. Consequently, we established a training set of 2,223 data and a validation set of 316 data.

**Figure 1.**
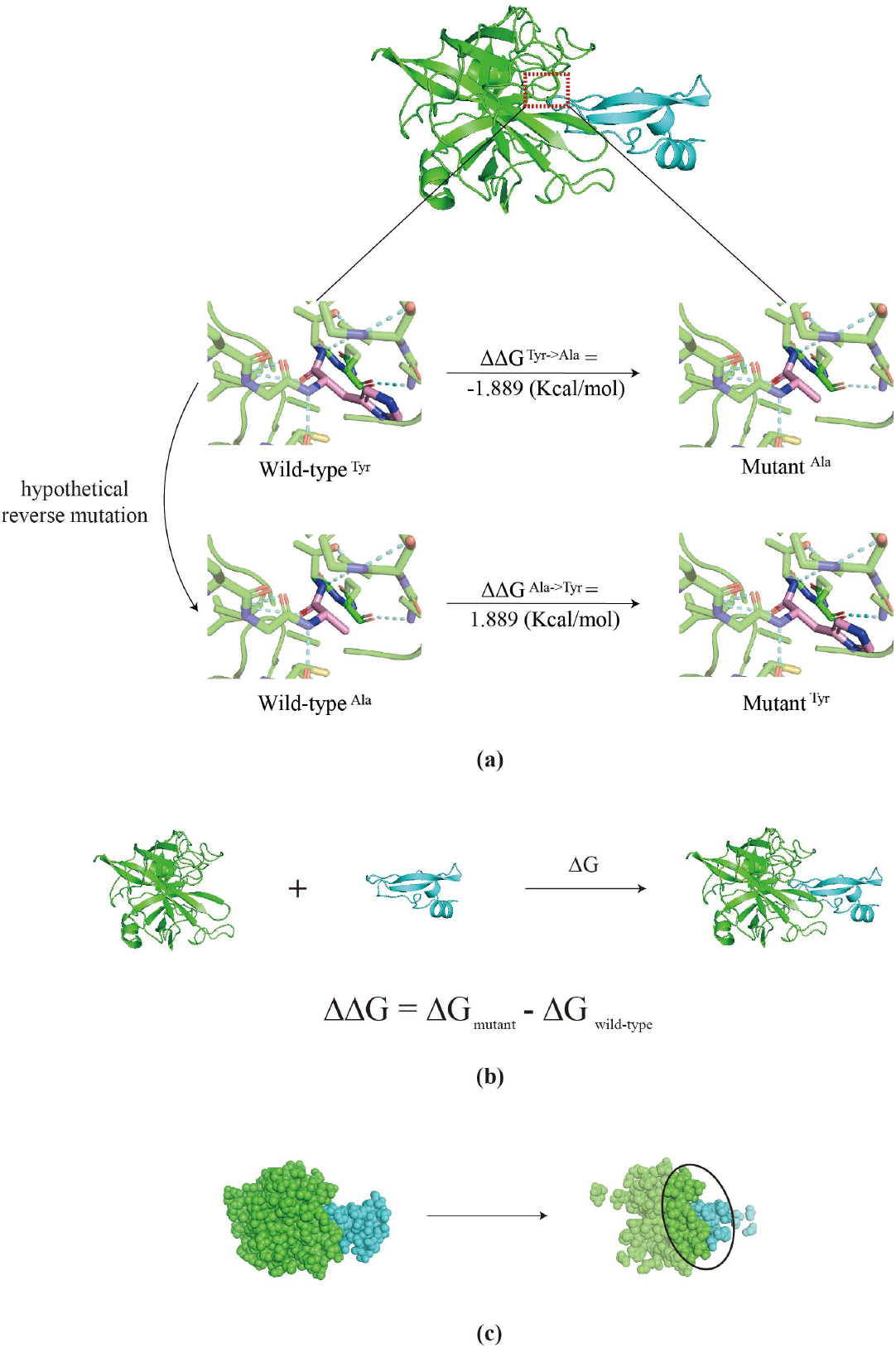
Data explanation and preprocessing. (a) One data includes two components: one is the experimental structure of the pre-mutation antibody-antigen complex, and the other is mutation information (mutation sites, original and mutated residues, and the resulting ΔΔ*G*). The hypothetical reverse mutation is introduced to balance the ΔΔ*G* distribution and augment the training data. (b) The prediction target ΔΔ*G* is defined by the difference in Δ? between the mutant and the original complex. Δ*G* is the change of Gibbs free energy that indicates the decrease in energy of an antibody-antigen interactions. (c) Residues in the interaction zone are identified and retained, while distal residues far from the interaction surface undergo coarse-graining, where multiple residues are aggregated into one.

### 2.2 Data Preprocessing

Residues close to the antibody-antigen interface contribute more to affinity, while those farther away less [34]. Therefore, during the preprocessing stage (**Figure 1c**), we categorize residues based on their distance from the contact surface and whether they are standard amino acids. Residues within 5.0 Å of the contact surface are designated as core residues, which will be identified and retained [35]. While residues beyond 5.0 Å are classified as peripheral residues that undergo coarse-graining, where multiple residues are aggregated into one [36]. Non-standard residues are considered invalid and are differentiated using a masking approach.

### 2.3 Structural Feature Extraction and Representation

We based our structural feature extraction and representation on the work of RefineGNN [36], representing the complex structure as a graph 𝒢 = 𝒱, ℰ with residues as nodes. The node features𝒱 = {*v*_1_, ⋯, *v*_2_} are defined by the dihedral angles 𝒟 of residues, while the edge features ℰ = {*e*_*ij*_|1 ≤ *i,j* ≤ *n, i* ≠ *j*} are characterized by sequence distance *E*_*pos*_, spatial distance *D*_*pos*_, and spatial rotation properties 𝒪.

#### Node Features Representation

For the representation of node features, we adhere to the Ramachandran plot [37], using 𝒟_*i*_ = ϕ_*i*_, ψ_*i*_, ω_*i*_ to denote the dihedral angle feature of the corresponding residues, which was more biological meaningful. The node features of residue *i* are computed by:

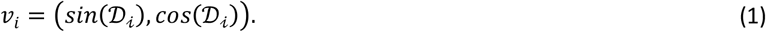

#### Edge Features Representation

To conserve computational resources and focus on key information, we chose the K-nearest neighbors to construct edge *e*_*ij*_, which represents the interactions between residue *i* and residue*j* :

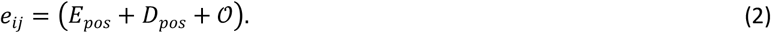

Sequence distance *E*_*pos*_ escribes the positional distance between residues connected by a peptide chain at the chain level, where sequence distance between chains holds no meaningful significance. Consequently, we did ablation analysis (see **Table S1**) to only consider the residue’s neighbors to its 30 adjacent upstream and downstream:

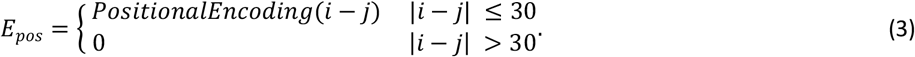

Spatial distance *D*_*pos*_ is calculated using the Euclidean distance between the *C*_α_ coordinates of residues, which is then transformed into a radial basis function representation [38]:

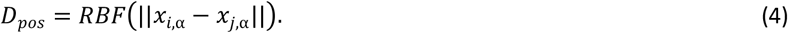

Spatial rotation properties 𝒪 are represented by a spatial rotation matrix in a quaternion function. The spatial rotation matrix 𝒪_*i*_ is calculated by an orientation matrix in a local coordinate system for each residue [39]:

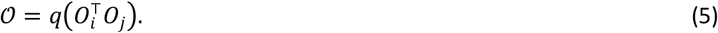

### 2.4 Sequential Feature Extraction and Progressive Encoding in Encoder

Sequential features include residue type *S*_*r*_ and context information *S*_*c*_. Residue types are encoded using an embedding layer, representing each residue as a corresponding vector. Context information is captured through a bidirectional gated recurrent unit (GRU) and weighted across sequence positions via an attention mechanism.

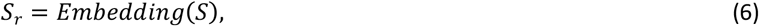

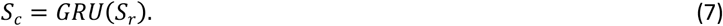

*S* is the indexed sequence, using number between 0 to 20 to represent each amino acid.

Afterwards, the Progressive Encoding (**Figure 2**) would integrate the above extracted features in three steps: Protein Structure Encoding (PSE), Residue-guided PSE (RPSE), and Context-embedded RPSE (CRPSE).

**Figure 2.**
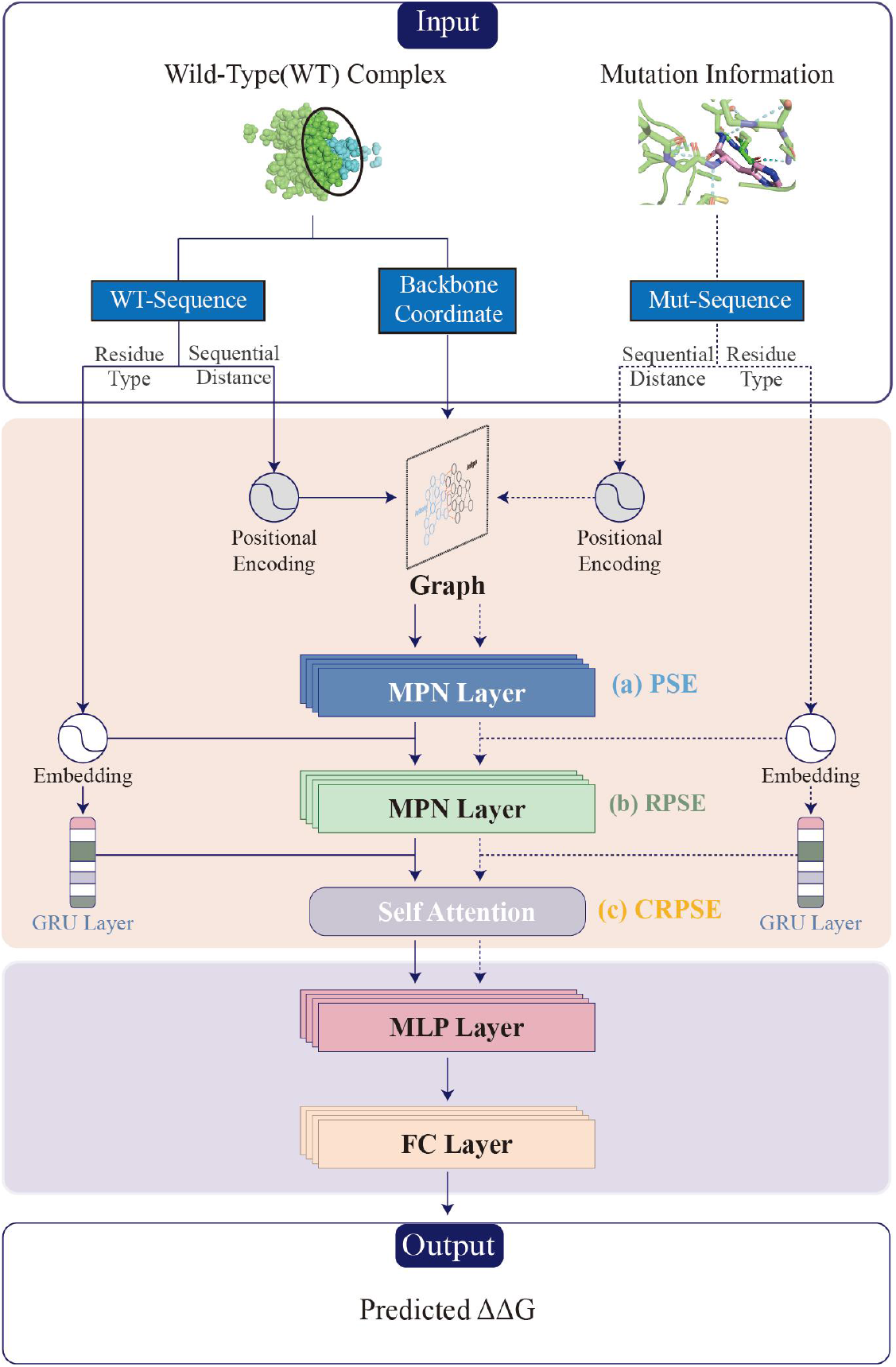
Workflow of abCAN. Model’s input includes two parts, one structure data of the wild-type antibody-antigen complex, and mutation information, including the type and location. Progressive Encoding involves the following steps: (a) Protein Structure Encoding (PSE): Structural features, including sequence distance, are first organized into a graph, which is then encoded using a message-passing network. (b) Residue-guided PSE (RPSE): On top of the structural encoding, residue type information is integrated using a message-passing network with different parameters. (c) Context-embedded RPSE: A GRU is used to extract sequence contextual information, with an attention mechanism applied to assign different weights to nodes at different sequence positions. After progressive encoding, two complex representations — one for the wild-type and one for the mutant — are obtained. These representations are passed through a MLP to learn the underlying residue interactions, followed by a fully connected layer to predict the final ΔΔ*G* value.

To be specific, the process begins with PSE that encodes structural features using a message-passing network (MPN) with parameter θ_1_. The initial graph is represented by the node features:

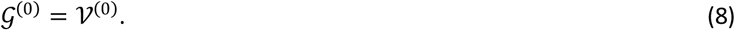

Then each layer *l*_θ_ in this 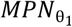 enrich the PSE with edge features ℰ:

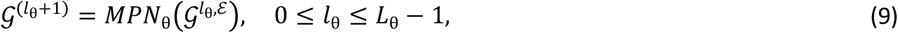

*L*_θ_ stands for the total number of layers in 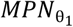.

Since this step does not differentiate between residue types, it treats pre- and post-mutation complexes with a shared PSE. Next, residue type information is introduced for each node in the encoded structural graph. This step distinguishes between the pre- and post-mutation complexes due to the presence of mutations in sequence. The implementation is carried out using another MPN parameterized by θ_2_. In the initial layers, only the residue type information of the corresponding nodes is considered. In the subsequent layers, the residue type of neighboring nodes is additionally incorporated, which are meant to further capture interactions between residues:

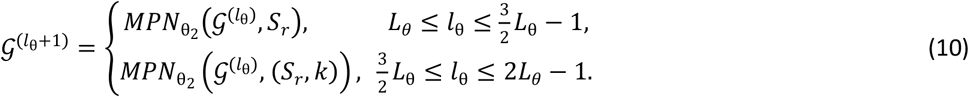

*k* represents the number of nearest neighbors selected based on the Euclidean distance.

Finally, sequence context information *S*_*c*_ is considered, where the attention mechanism assigns additional weight to residue nodes near the interaction region:

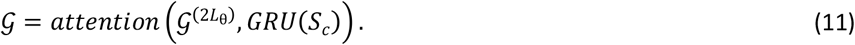

Each antibody-antigen complex has a corresponding graph representation 𝒢, a paired representation (𝒢_*wild*_*types*,_𝒢_*mutant*_) would then be input into the corresponding network for learning.

### 2.5 Predictor Architecture

#### Multi-Layer Perceptron for Learning Complex Representations

The paired pre- and post-mutation complex representation (𝒢_*wild*_*types*,_𝒢_*mutant*_) is fed into a multi-layer perceptron (MLP) to learn the interaction features between the antigen and antibody within the complex.

#### Fully Connected Layer for Predicting Affinity Change (ΔΔ*G*)

The interaction features learned in the previous step are used to compute the difference between pre- and post-mutation complexes, which is then fed into a fully connected layer to predict the change in affinity (ΔΔ*G*) for each mutant complex.

#### Loss Calculation and Backpropagation

The loss ℒ is defined as the mean squared error between the predicted values and the actual values. Adam optimizer is used to perform backpropagation, completing one iteration of model training.

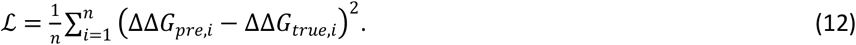

### 2.6 Model Evaluation Metrics and Methods

We selected Root Mean Square Error (RMSE) and Pearson Correlation Coefficient (PCC) as two evaluation metrics, which respectively measure the accuracy and precision of the model’s predictions. PCC is calculated using the SciPy library in Python, while RMSE is calculated using the formula below:

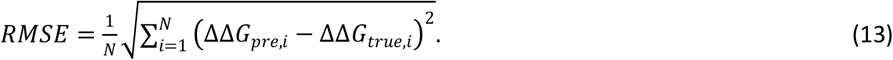

We performed k-fold cross-validation on training and validation data to assess the model’s robustness and eliminate the impact of different training set partitions on model performance. To compare the performance of abCAN with other SOTA methods, we use benchmark M305 to evaluate prediction accuracy (in terms of RMSE and PCC).

Additionally, to explore the contribution of each feature component, we conducted ablation analysis. All the above implementations were conducted using the PyTorch framework (version 2.0.0), on an Ubuntu 22.04.04 LTS system with an Intel Xeon w7-3445 CPU and a single NVIDIA GeForce RTX 4090 GPU. The specific results will be discussed in the Results section.

## 3 Results

### 3.1 Model Performance

We trained the model using 2,223 training data, with an initial learning rate set to 0.001. The learning rate was reduced by 15% if there was no convergence after 10 epochs. The optimal hyperparameter combination for abCAN was determined through a random search on the training and validation sets (see **Table S2**). The final model achieved a RMSE of 1.195 kcal/mol^-1^ and a PCC of 0.841 (p-value < 10^−65^) on the benchmark M305, the illustration of prediction can be seen on **Figure 3**.

**Figure 3.**
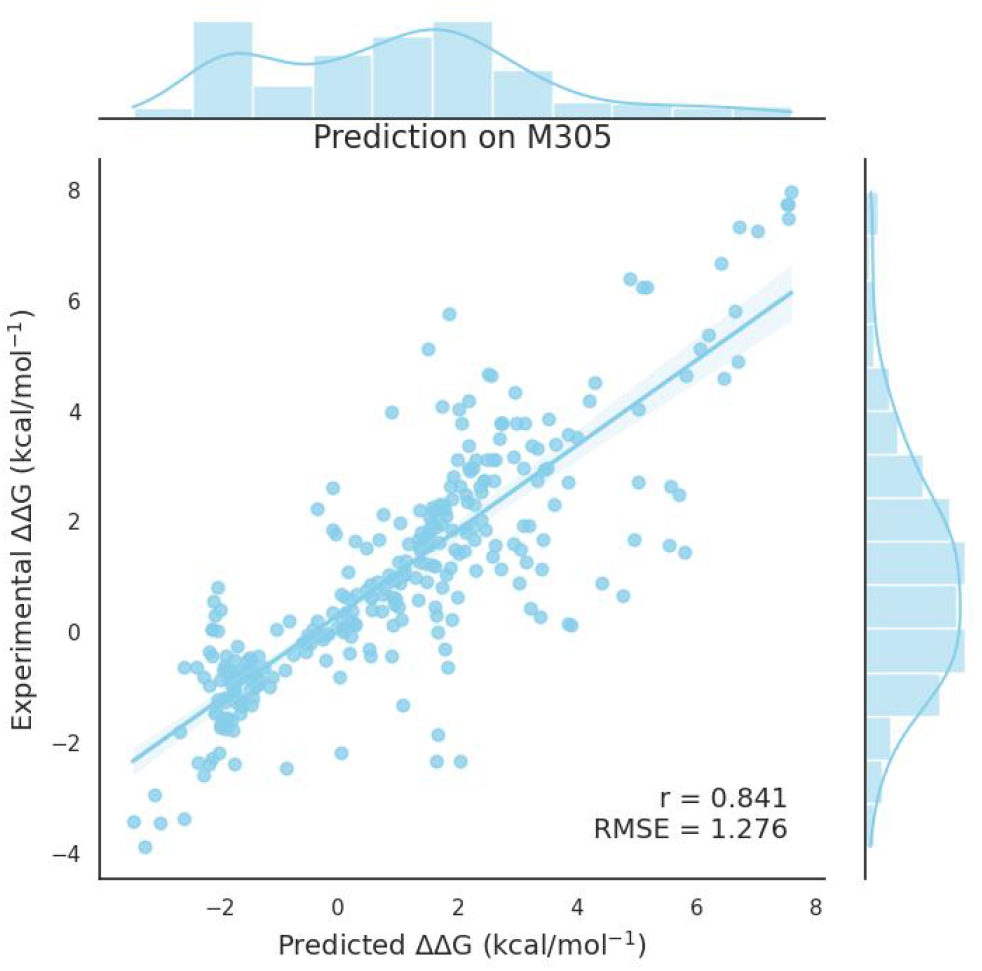
ΔΔ*G* predicted by abCAN. Shown is the prediction against the experimental determination on benchmark M305. P-value of the PCC is less than 10^−23^.

### 3.2 Performance Comparison on Benchmark Dataset

Table 1. summarizes the performance metrics of various methods (MutaBind2 [40], DDG predictor [41], GeoPPI [30]) on benchmark M305, with abCAN displaying outstanding performance surpassing other models, even though MutaBind2 had prior exposure to some data from benchmark M305. Notably, unlike other methods, we didn’t differentiate between the number of mutations during abCAN’s training process; yet, it demonstrates excellent predictive accuracy. This result suggests that our novel architecture, Progressive Encoding, enables the model to learn the underlying principles of antibody-antigen interactions. We provide a detailed analysis of one single and one double mutations example (see **Figure 4**) to illustrate abCAN’s function. The changes in the affinity actually suggest the changes of the bonding in the complex. For complex 1EAW (see **Figure 4a**), the original histidine is hydrophilic and possesses an aromatic side chain. After mutating to alanine, the region becomes more hydrophobic, and the steric hindrance is significantly reduced, leading to a change in the local conformation. Consequently, one additional hydrogen bond is formed, enhancing the overall affinity. As for complex 2B2X that consists of two mutations, the changing force is not hydrogen bond but π-π stacking interaction between tyrosine residues in position 48 and 52 of the origin L chain [42]. Such π-π stacking interaction significantly contributes to the structural stability of the complex by enhancing non-covalent binding energy, whereas mutations disrupt it, leading to an increase in ΔΔ*G*. The mutation on 53rd residue actually brings in a new hydrogen bond between 52nd residue and 48th residue, which may explain the large negative prediction made by DDG predictor. This is an advanced prediction example since normally it is hydrogen bond that significantly contributes to protein-protein interaction [43], but abCAN can still successfully identify the overall binding changes, indicating its ability of capturing complicated interaction patterns. abCAN spent the minimum inference speed as well, and the complete comparison of inference speed and occupation can be found in **Table S3**.

### 3.3 K-fold and Split-by-Structure Cross-validation

To validate the robustness of the model, we conducted 5-fold, 10-fold, and split-by-structure cross-validation (SSCV) [30]. The performance of 5-fold and 10-fold cross-validation is stable, with no outliers. The average PCC is 0.710 and 0.741, while the average RMSE is 1.296 kcal/mol^-1^ and 1.232 kcal/mol^-1^, respectively. The complete results can be found in **Table S4** and **Table S5**.

**Table 1:** Comparison of Performance on benchmark M305. † The results of MutaBind2 were obtained from its publicly available web server. However, since its training data partially overlaps with the benchmark M305, the reported metrics may be deceptive better. ‡ The results were obtained based on the released source codes.

| Method | PCC | RMSE (kcal/mol) |
| --- | --- | --- |
| abCAN | 0.841 | 1.195 |
| MutaBind2† | 0.817 | 1.273 |
| DDG predictor‡ | 0.469 | 3.680 |
| GeoPPI‡ | 0.368 | 2.119 |

**Figure 4.**
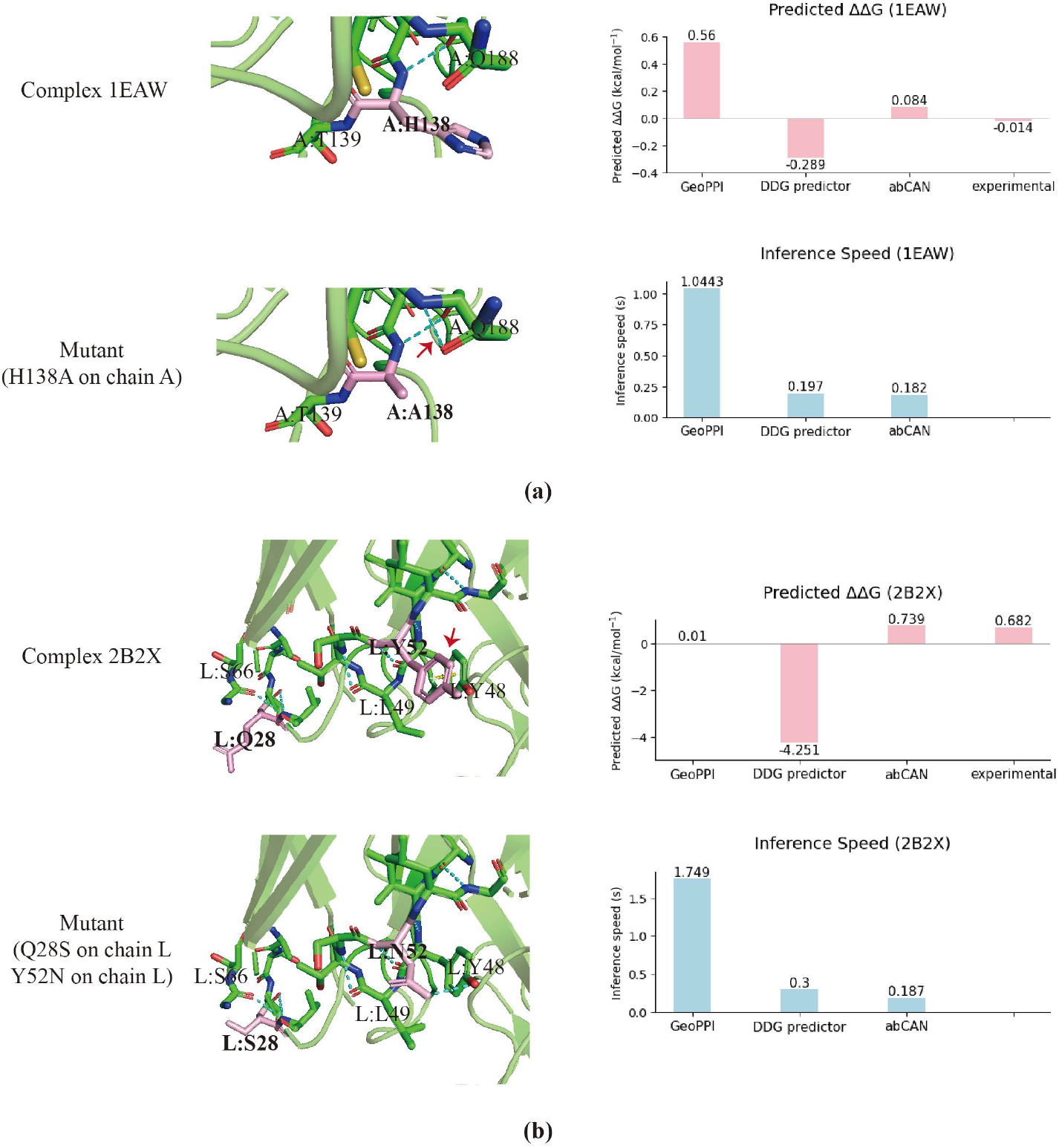
ΔΔ*G* predicted by abCAN corresponds to the binding changes. Left panel displays the local conformation difference, where the form or loss of non-covalent interaction would be indicated by red arrows. The right panel illustrates the comparison of ΔΔ*G* value as well as inference speed of each model. (a) Binding changes of complex 1EAW before and after the mutation from histidine (H) to alanine (A) at the 23rd residue of chain A, leading to an extra hydrogen bond. (b) Binding changes of complex 2B2X before and after two mutations, one from glutamine (Q) to serine (S) at the 28th residue of chain L, and the other from tyrosine (Y) to asparagine (N) at the 52nd residue of chain L, disrupting the π-π stacking interaction between the original two tyrosine residues.

In the SSCV (see **Figure 5**), we evaluated performance by leaving out data corresponding to one structure from the benchmark M305 at a time. The aim is to see whether the model’s performance is influenced by specific structures, since some structures have more mutation data and might dominate the performance. For each comparing method, a set of 34 PCCs was obtained, and we performed pairwise comparisons between abCAN and the other two using the Wilcoxon signed-rank test. GeoPPI exhibits fewer outliers, but the overall PCC distribution is below 0.4. DDG predictor shows a higher PCC distribution between 0.4 and 0.5, though it has a higher number of outliers, which affects model stability. In contrast, abCAN demonstrates fewer outliers and a more concentrated PCC distribution, with values above 0.85. Furthermore, the results of Wilcoxon signed-rank test revealed significant differences, as both W=0.0 and p-value are less than 10^−10^. Consequently, we conclude that abCAN significantly outperforms the other models in terms of stability and accuracy.

**Figure 5.**
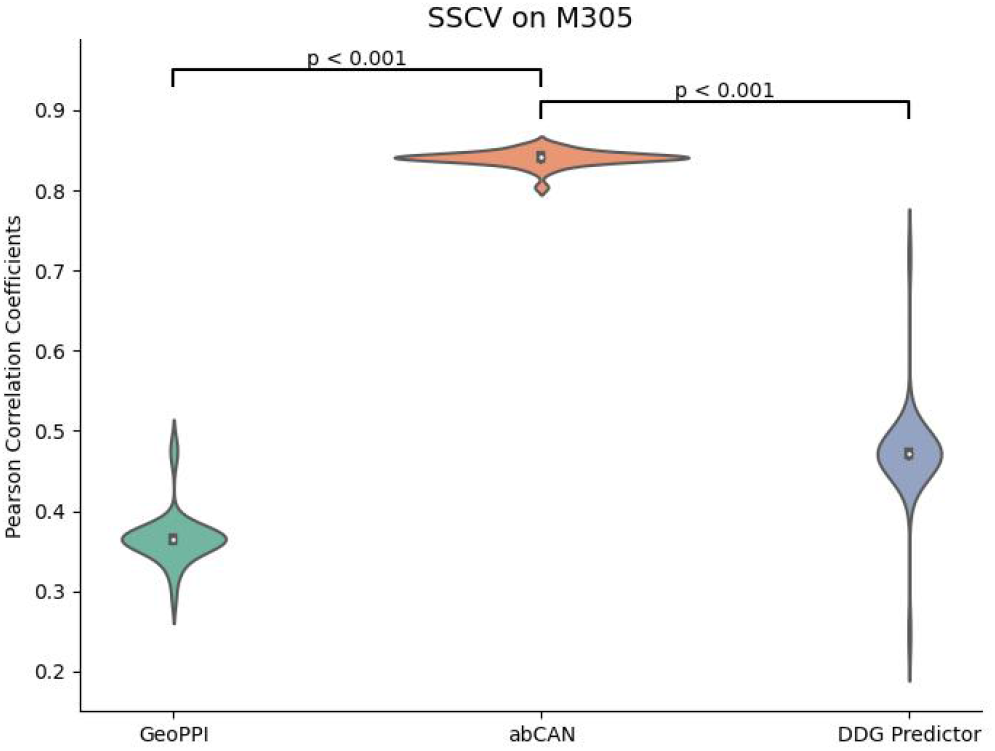
Distribution of Pearson Correlation Coefficient (PCC) obtained by different models in SSCV. Each violin is composed of 34 PCCs, with each PCC value representing the testing result after excluding one structure from the benchmark M305. The p-values marked between each violin are the results of the Wilcoxon signed-rank test, and both values are less than 0.05, indicating a significantly improved performance of abCAN.

### 3.4 Ablation analysis

To assess the contribution of each feature encoding module to the model’s predictive performance, we conducted ablation analysis. With the hyperparameter combination fixed, we retrained the model by removing one encoding module at a time and compared the performance on the benchmark M305. The specific results are presented in **Table 2**.

**Table 2:** Ablation analysis of different encoding modules. **Progressive Encoding:** our novel way of encoding structural, residues and contextual features in a progressive manner.

| Feature | M305 |  |
| --- | --- | --- |
|  | PCC | RMSE (kcal/mol) |
| - | 0.841 | 1.195 |
| interface attention | 0.760 | 1.507 |
| node dihedral angle | 0.748 | 1.484 |
| spatial distance | 0.738 | 1.467 |
| sequence distance | 0.720 | 1.963 |
| rotation direction | 0.693 | 1.579 |
| sequence attention | 0.041 | 2.213 |
| Progressive Encoding | 0.036 | 2.025 |

As shown in **Table 2**, excluding any module results in a decline in the model’s performance. Modules such as interface attention, node dihedral angle, spatial distance, sequence distance and rotation direction represent specific biological features extracted from structural or sequential data, each contributing to the accuracy (approximately 0.3 RMSE) and precision (around 0.1 PCC). Notably, excluding sequence distance leads to a high RMSE of 1.963 kcal/mol^-1^, highlighting the equal or even greater importance of sequential distance compared to spatial distance. Although each feature is effective, none would be as impactful without Progressive Encoding, which carefully integrates these features using the most suitable network architectures. The exclusion of Progressive Encoding results in the largest drop in PCC, with the model failing to converge, which is an issue not observed with other modules. This suggests abCAN’s significant potential for capturing intrinsic patterns within biological macromolecules like antibody-antigen.

Furthermore, the sequence attention mechanism, designed to capture the contextual information within the complex sequence, also plays a vital role. Its exclusion corresponds to the highest RMSE, further illustrating the strong relationship between sequential information and predictive accuracy.

## 4 Discussion

Antibody affinity prediction is a pivotal step in antibody optimization, significantly advancing immunotherapy, scientific discovery, and disease prevention. Machine learning has demonstrated promising applications across various domains, including computer vision, natural language processing [44], and small molecule modeling. However, in the context of protein macromolecules such as antibodies and antigens, the challenge lies in developing models capable of learning complex interactions between chains in a three-dimensional space [45]. The limited availability of mutant structure data in this area, compared to the aforementioned fields, further constrains the development of deep learning in antibody macromolecule research. A prevailing consensus among mainstream approaches is to predict mutant structures and encode antibody-antigen complexes as graphs, where residues or atoms are treated as nodes and interaction forces as edges [46]. A representative example is GeoPPI [30], which uses atoms as nodes and assigns an edge between atoms within a 3 Å distance; node features include atom type, residue type, chain name, interaction interface status, and three-dimensional coordinates. Although these methods provide the model with a comprehensive set of information, the nuanced distinction between information is not understandable to models, as all features are concatenated in a relatively simple manner. We believe that the key to breakthroughs lies in systematically integrating biological knowledge with appropriate neural network architectures [47].

Therefore, we propose two innovations: Progressive Encoding and interface attention. Progressive Encoding separates three types of features (structural information, residue types, and sequence context) and employs suitable neural network modules for encoding before systematically integrating them in a hierarchical manner. In this encoding approach, the protein structure encoding (PSE) serves as the foundation, delineating the framework for graph encoding, which is characterized by its insensitivity to local mutations [48]. Residue types provide local details at the sequence level, assigning identity significance to each graph node. Finally, sequence context information determines the ultimate representation from the perspective of protein language. Clearly, we place greater emphasis on the latter sequential information, as mutant sequences can be accurately generated; however, large-scale experimental determination of mutant structures is currently not feasible [19]. Although numerous algorithmic tools for structure prediction exist, they may introduce conformational differences in the mutated structure compared to the original experimental structure, as well as inherent prediction errors [49].

Progressive Encoding effectively circumvents the use of mutant structures, enabling the model to make predictions based on robust experimental data. Consequently, abCAN requires only the structure of the pre-mutant complex and mutation information as input, aligning more closely with the practical process of antibody mutation optimization in research. Additionally, the benchmark M305 contains a higher proportion of multiple mutation data compared to single ones, which introduces greater structural perturbations. abCAN demonstrates superior performance under these conditions, supporting the concept of Progressive Encoding, emphasizing sequence-level information based on backbone structure.

Additionally, the interface attention enhances the general sequence attention by computing extra weights for the antibody-antigen interaction region, enabling the model to focus more on this critical area. Despite that antibody-antigen interactions represent a specialized subset of protein-protein interactions, antigen themselves are general proteins without conservative structures like antibodies. The architecture of abCAN is fully transferable to the representation of individual proteins or protein-protein interactions. We plan to further explore this potential in future work.

However, the premise of sharing PSE is that limited number of mutations won’t influence the backbone structure. We would also incorporate predicted complex structures in our feature work with the help of Alphafold3 [50]. We anticipate that abCAN, with its more practical input and shorter prediction time, will become a powerful tool in antibody optimization, aiding in the development of antibody therapeutics.

## 5 Supplementary Information

**Table S1** To result of ablation analysis to determine the value of *k*_*seq*_. The results indicate that value 30 contributes to the optimal performance.

**Table S2** The best hyperparameters combination selected through random search approach over *L*_θ_ ∈ {4,6,8}, *C*∈ {8,16,32}, *N*_*r*_ ∈ {8,16,32}, *P*_*pc*_ ∈ {16,32,64}, *B*∈ {8,16,32}, *d* ∈ {0.1,0.3}, *D* ∈ {128,256} on the validation set.

**Table S3** illustrates the time and CPU, GPU memory usage required to predict all samples in the benchmark M305. abCAN exhibited a significant reduction in prediction time.

**Table S4** and **Table S5** are complete results of 5-fold and 10-fold cross-validation.

**Figure S1** is the visualization of prediction in **Table 1**.

## 6 Acknowledgement

This work was supported in part by the National Natural Science Foundation of China under grant 62401272, the Talent Start-up Funding Project of Nanjing University of Information Science and Technology under grant 2024r090, the Program from Anhui Key Laboratory of Computational Medicine and Intelligent Health under grant AHCM2024Z001, and by the Noncommunicable Chronic Diseases-National Science and Technology Major Project.

## References

[1] L. Qian et al., “The dawn of a new era: targeting the “undruggables” with antibody-based therapeutics,” Chemical reviews, vol. 123, no. 12, pp. 7782–7853, 2023.

[2] H.-P. Peng et al., “Antibody CDR amino acids underlying the functionality of antibody repertoires in recognizing diverse protein antigens,” Scientific Reports, vol. 12, no. 1, p. 12555, 2022.

[3] K. Tsuchikama, Y. Anami, S. Y. Ha, and C. M. Yamazaki, “Exploring the next generation of antibody–drug conjugates,” Nature Reviews Clinical Oncology, vol. 21, no. 3, pp. 203–223, 2024.

[4] B. G. De la Torre and F. Albericio, “The pharmaceutical industry in 2023: An analysis of FDA drug approvals from the perspective of molecules,” Molecules, vol. 29, no. 3, p. 585, 2024.

[5] J. Li et al., “Affinity maturation of antibody fragments: A review encompassing the development from random approaches to computational rational optimization,” International journal of biological macromolecules, p. 125733, 2023.

[6] A. Vangone and A. M. Bonvin, “Contacts-based prediction of binding affinity in protein–protein complexes,” elife, vol. 4, p. e07454, 2015.

[7] R. Singh, P. Chandley, and S. Rohatgi, “Recent Advances in the Development of Monoclonal Antibodies and Next-Generation Antibodies,” ImmunoHorizons, vol. 7, no. 12, pp. 886–897, 2023, doi: 10.4049/immunohorizons.2300102.

[8] C. Geng, L. C. Xue, J. Roel-Touris, and A. M. Bonvin, “Finding the ΔΔG spot: Are predictors of binding affinity changes upon mutations in protein–protein interactions ready for it?,” Wiley Interdisciplinary Reviews: Computational Molecular Science, vol. 9, no. 5, p. e1410, 2019.

[9] R. Khetan et al., “Current advances in biopharmaceutical informatics: guidelines, impact and challenges in the computational developability assessment of antibody therapeutics,” in MAbs, 2022, vol. 14, no. 1: Taylor & Francis, p. 2020082.

[10] S. Izadi, T. W. Patapoff, and B. T. Walters, “Multiscale coarse-grained approach to investigate self-association of antibodies,” Biophysical journal, vol. 118, no. 11, pp. 2741–2754, 2020.

[11] T. Sulea, V. Vivcharuk, C. R. Corbeil, C. Deprez, and E. O. Purisima, “Assessment of solvated interaction energy function for ranking antibody–antigen binding affinities,” Journal of chemical information and modeling, vol. 56, no. 7, pp. 1292–1303, 2016.

[12] A. Tennenhouse et al., “Computational optimization of antibody humanness and stability by systematic energy-based ranking,” Nature biomedical engineering, pp. 1–15, 2023.

[13] Y. Kurumida, Y. Saito, and T. Kameda, “Predicting antibody affinity changes upon mutations by combining multiple predictors,” Scientific reports, vol. 10, no. 1, p. 19533, 2020.

[14] E. K. Makowski et al., “Co-optimization of therapeutic antibody affinity and specificity using machine learning models that generalize to novel mutational space,” Nature communications, vol. 13, no. 1, p. 3788, 2022.

[15] W. Wilman et al., “Machine-designed biotherapeutics: opportunities, feasibility and advantages of deep learning in computational antibody discovery,” Briefings in Bioinformatics, vol. 23, no. 4, p. bbac267, 2022.

[16] P. Tessier, “Simplifying Complex Antibody Engineering Using Machine Learning,” in 2022 AIChE Annual Meeting, 2022: AIChE.

[17] T. Clark et al., “Machine Learning-Guided Antibody Engineering That Leverages Domain Knowledge To Overcome The Small Data Problem,” bioRxiv, p. 2023.06. 02.543458, 2023.

[18] Y. Zhu, W. Li, and T. Li, “A hybrid artificial immune optimization for high-dimensional feature selection,” Knowledge-Based Systems, vol. 260, p. 110111, 2023.

[19] J. Dunbar et al., “SAbDab: the structural antibody database,” Nucleic acids research, vol. 42, no. D1, pp. D1140–D1146, 2014.

[20] J. Parkinson, R. Hard, and W. Wang, “The RESP AI model accelerates the identification of tight-binding antibodies,” Nature Communications, vol. 14, no. 1, p. 454, 2023.

[21] L. Li, E. Gupta, J. Spaeth, L. Shing, T. Bepler, and R. S. Caceres, “Antibody representation learning for drug discovery,” arXiv preprint arXiv:2210.02881, 2022.

[22] E. Davidson and B. J. Doranz, “A high-throughput shotgun mutagenesis approach to mapping B-cell antibody epitopes,” Immunology, vol. 143, no. 1, pp. 13–20, 2014.

[23] W. Jin, R. Barzilay, and T. Jaakkola, “Antibody-antigen docking and design via hierarchical structure refinement,” in International Conference on Machine Learning, 2022: PMLR, pp. 10217–10227.

[24] B. K. Rai, J. R. Apgar, and E. M. Bennett, “Low-data interpretable deep learning prediction of antibody viscosity using a biophysically meaningful representation,” Scientific Reports, vol. 13, no. 1, p. 2917, 2023.

[25] N. L. Miller, T. Clark, R. Raman, and R. Sasisekharan, “Learned features of antibody-antigen binding affinity,” Frontiers in Molecular Biosciences, vol. 10, p. 1112738, 2023.

[26] S. Sirin, J. R. Apgar, E. M. Bennett, and A. E. Keating, “AB-bind: antibody binding mutational database for computational affinity predictions,” Protein Science, vol. 25, no. 2, pp. 393–409, 2016.

[27] J. Jankauskaitė, B. Jiménez-García, J. Dapkūnas, J. Fernández-Recio, and I. H. Moal, “SKEMPI 2.0: an updated benchmark of changes in protein–protein binding energy, kinetics and thermodynamics upon mutation,” Bioinformatics, vol. 35, no. 3, pp. 462–469, 2019.

[28] I. H. Moal and J. Fernández-Recio, “SKEMPI: a structural kinetic and energetic database of mutant protein interactions and its use in empirical models,” Bioinformatics, vol. 28, no. 20, pp. 2600–2607, 2012.

[29] Y. Myung, C. H. Rodrigues, D. B. Ascher, and D. E. Pires, “mCSM-AB2: guiding rational antibody design using graph-based signatures,” Bioinformatics, vol. 36, no. 5, pp. 1453–1459, 2020.

[30] X. Liu, Y. Luo, P. Li, S. Song, and J. Peng, “Deep geometric representations for modeling effects of mutations on protein-protein binding affinity,” PLoS computational biology, vol. 17, no. 8, p. e1009284, 2021.

[31] G. Thiltgen and R. A. Goldstein, “Assessing predictors of changes in protein stability upon mutation using self-consistency,” PloS one, vol. 7, no. 10, p. e46084, 2012.

[32] D. E. Pires and D. B. Ascher, “mCSM-AB: a web server for predicting antibody–antigen affinity changes upon mutation with graph-based signatures,” Nucleic acids research, vol. 44, no. W1, pp. W469–W473, 2016.

[33] C. H. Rodrigues, D. E. Pires, and D. B. Ascher, “DynaMut: predicting the impact of mutations on protein conformation, flexibility and stability,” Nucleic acids research, vol. 46, no. W1, pp. W350–W355, 2018.

[34] T. Ramaraj, T. Angel, E. A. Dratz, A. J. Jesaitis, and B. Mumey, “Antigen–antibody interface properties: Composition, residue interactions, and features of 53 non-redundant structures,” Biochimica et Biophysica Acta (BBA)-Proteins and Proteomics, vol. 1824, no. 3, pp. 520–532, 2012.

[35] M. Wang, D. Zhu, J. Zhu, R. Nussinov, and B. Ma, “Local and global anatomy of antibody-protein antigen recognition,” Journal of Molecular Recognition, vol. 31, no. 5, p. e2693, 2018.

[36] W. Jin, J. Wohlwend, R. Barzilay, and T. Jaakkola, “Iterative refinement graph neural network for antibody sequence-structure co-design,” arXiv preprint arXiv:2110.04624, 2021.

[37] G N R Ramachandran, C.; Sasisekharan, V., “Stereochemistry of polypeptide chain configurations.”,” J. mol. Biol, vol. 7, pp. 95–99, 1963.

[38] J. Qiao, F. Li, C. Yang, W. Li, and K. Gu, “A self-organizing RBF neural network based on distance concentration immune algorithm,” IEEE/CAA Journal of Automatica Sinica, vol. 7, no. 1, pp. 276–291, 2019.

[39] J. Ingraham, V. Garg, R. Barzilay, and T. Jaakkola, “Generative models for graph-based protein design,” Advances in neural information processing systems, vol. 32, 2019.

[40] N. Zhang et al., “MutaBind2: predicting the impacts of single and multiple mutations on protein-protein interactions,” Iscience, vol. 23, no. 3, 2020.

[41] S. Shan et al., “Deep learning guided optimization of human antibody against SARS-CoV-2 variants with broad neutralization,” Proceedings of the National Academy of Sciences, vol. 119, no. 11, p. e2122954119, 2022.

[42] Y. Zhao et al., “Conformational preferences of π–π stacking between ligand and protein, analysis derived from crystal structure data geometric preference of π–π interaction,” Interdisciplinary Sciences: Computational Life Sciences, vol. 7, pp. 211–220, 2015.

[43] C. N. Pace et al., “Contribution of hydrogen bonds to protein stability,” Protein Science, vol. 23, no. 5, pp. 652–661, 2014.

[44] M.-W. C. Jacob Devlin, Kenton Lee, Kristina Toutanova, “BERT: Pre-training of Deep Bidirectional Transformers for Language Understanding,” Proceedings ofthe 2019 Conference ofthe Association for Computational Linguistics, pp. 4171–4186, 2019.

[45] M. K. Goshisht, “Machine learning and deep learning in synthetic biology: Key architectures, applications, and challenges,” ACS omega, vol. 9, no. 9, pp. 9921–9945, 2024.

[46] M. Xu et al., “Graph Neural Networks for Protein-Protein Interactions-A Short Survey,” arXiv preprint arXiv:2404.10450, 2024.

[47] A. Prosz et al., “Biologically informed deep learning for explainable epigenetic clocks,” Scientific Reports, vol. 14, no. 1, p. 1306, 2024.

[48] A. A. Rosenberg, N. Yehishalom, A. Marx, and A. M. Bronstein, “An amino-domino model described by a cross-peptide-bond Ramachandran plot defines amino acid pairs as local structural units,” Proceedings of the National Academy of Sciences, vol. 120, no. 44, p. e2301064120, 2023.

[49] M. L. Fernández-Quintero et al., “Challenges in antibody structure prediction,” in MAbs, 2023, vol. 15, no. 1: Taylor & Francis, p. 2175319.

[50] J. Abramson et al., “Accurate structure prediction of biomolecular interactions with AlphaFold 3,” Nature, pp. 1–3, 2024.

