## Supplementary figures and images for "abCAN: a Practical and Novel Attention Network for Predicting Mutant Antibody Affinity"

### Figure S1

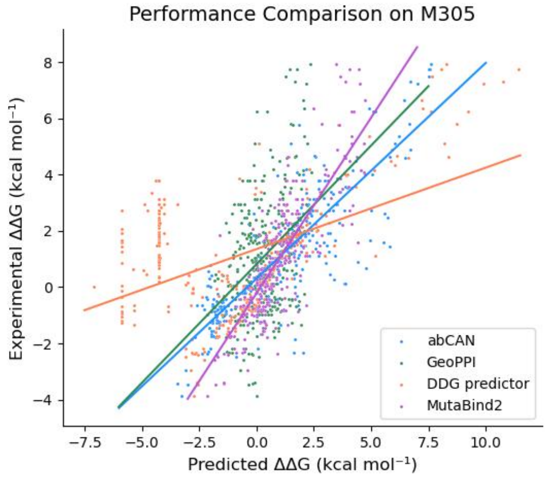


**Figure S1. Comparison of predicting performance of different models on M305.**
