## Supplementary material for "abCAN: a Practical and Novel Attention Network for Predicting Mutant Antibody Affinity": Table S1

**Table S1: Ablation analysis of $K_{seq}$ value.** $K_{seq}$: the number of upstream and downstream neighbors in the sequence positional encoding.

| $K_{seq}$ | PCC | RMSE (kcal/mol) |
|---|---|---|
| **10** | 0.791 | 1.316 |
| **20** | 0.730 | 1.559 |
| **30** | 0.841 | 1.195 |
| **40** | 0.745 | 1.648 |
