## Supplementary material for "abCAN: a Practical and Novel Attention Network for Predicting Mutant Antibody Affinity": Table S2

**Table S2: Selected Hyperparameters of abCAN.** $L_\theta$: depth in both MPN modules, $N_c$: number of residues in one coarse-grained module, $N_r$: number of central points of the radial basis function, $D_{pc}$: hidden dimensions in sequence positional encoding, $B$: batch size, $d$: dropout rate, $D$: hidden size in neural networks.

| Hyperparameter | Selected values |
|---|---|
| $L_\theta$ | 4 |
| $N_c$ | 32 |
| $N_r$ | 8 |
| $D_{pc}$ | 16 |
| $B$ | 8 |
| $d$ | 0.1 |
| $D$ | 128 |
