## Supplementary material for "abCAN: a Practical and Novel Attention Network for Predicting Mutant Antibody Affinity": Table S3

**Table S3: Complete results of 5-fold cross-validation. Valid fold:** the fold that was used as the validation set in each split.

| Valid fold | PCC | RMSE (kcal/mol) |
|---|---|---|
| **0** | 0.756 | 1.323 |
| **1** | 0.688 | 1.393 |
| **2** | 0.750 | 1.104 |
| **3** | 0.702 | 1.343 |
| **4** | 0.652 | 1.319 |
