## Supplementary material for "abCAN: a Practical and Novel Attention Network for Predicting Mutant Antibody Affinity": Table S4

**Table S4: Complete results of 10-fold cross-validation. Valid fold:** the fold that was used as the validation set in each split.

| Valid fold | PCC | RMSE (kcal/mol) |
|---|---|---|
| **0** | 0.695 | 1.290 |
| **1** | 0.789 | 1.006 |
| **2** | 0.667 | 1.458 |
| **3** | 0.756 | 1.415 |
| **4** | 0.802 | 1.159 |
| **5** | 0.681 | 1.057 |
| **6** | 0.764 | 1.151 |
| **7** | 0.781 | 1.158 |
| **8** | 0.773 | 1.191 |
| **9** | 0.702 | 1.430 |
