## Supplementary material for "abCAN: a Practical and Novel Attention Network for Predicting Mutant Antibody Affinity": Table S5

**Table S5: Comparison of inference time and memory usage on M305.**

| Method | Inference time (s) | CPU memory (MiB) | GPU memory (MiB) |
|---|---|---|---|
| **abCAN** | 4.170 | 1024 | 712 |
| **DDG predictor** | 53.693 | 752.4 | 550 |
| **GeoPPI** | 651.93 | 1331.2 | 15 |
